# Evaluating a Standard Benchmark for Gene Prioritization: The InheriNext® Algorithm’s Integration of Genomic and Phenotypic Information

**DOI:** 10.1101/2025.02.25.640147

**Authors:** Ju-Yuan Chang, Kuan-Tsung Li, Yu-Shen Tsai, Michael Kubal, Aaron M. Hamby, Naomi Thomson, Jonathan Sheridan, Shiloh Barfield, Randy Rutz, Frank S. Ong, Ramon Felciano, Scott Kahn, Shao-Min Wu

## Abstract

This study presents a comprehensive benchmark analysis of InheriNext^®^, a domain-specific, AI-powered tool designed for phenotype-driven pathogenic variant prioritization. For this study, 7,244 whole exome test cases were generated using phenotype and genotype data from Phenopackets, along with pools of variants from healthy individuals to serve as genomic backgrounds. Performance was evaluated across diverse testing scenarios and compared against four established tools. The results show InheriNext^®^ achieving a 98.6% sensitivity in identifying pathogenic variants and consistent performance across diverse tests for variant types, phenotype counts, and disease groups—supporting the robustness and adaptability of its methodology. Sharing these benchmarking results and samples is intended to drive progress by assisting clinicians and researchers in evaluating interpretation tools and identifying areas for improvement.

## Introduction

Rare diseases, defined as conditions affecting fewer than 200,000 individuals in the United States or fewer than 1 in 2,000 people in the European Union, still collectively impact over 400 million people worldwide, highlighting a significant global health challenge [1]. Many of these diseases, often caused by subtle genetic mutations, go undiagnosed for extended periods. Accurate identification of genetic variants is essential; however, with over 10 million genetic variations present in an average human genome, effective prioritization becomes crucial. Advanced sequencing technologies, such as Whole Exome Sequencing (WES) and Whole Genome Sequencing (WGS), provide efficient methods for profiling genetic data. Through these sequencing methods, a patient’s specific genetic variants are identified, followed by prioritization based on the pathogenicity of the variants and the phenotypes relevant to the patient [2]. Previous research has highlighted the benefits of enhancing diagnostic accuracy by integrating phenotype data into variant prioritization algorithms [3, 4]. However, there remains significant room for improvement. For example, some consequences of genetic variants are difficult to interpret, as their functional impacts may not correspond to their actual pathogenicity.

Several current computational tools utilize clinical phenotype data annotated with HPO terms to rank candidate genes based on established phenotypic and genetic knowledge. Exomiser employs a logistic regression model that integrates variant-based and gene-based scores to generate a final prioritization score [5]. Variant-based scores are influenced by allele frequency, variant type, and pathogenicity predictions from tools. LIRICAL adopts a Bayesian statistical framework to assess candidate diagnoses by computing posterior probabilities based on likelihood ratios (LRs). It integrates *in silico* pathogenicity predictions and phenotype-based LRs, refining genotype-disease associations and improving diagnostic accuracy, particularly in rare disease contexts [6]. Xrare utilizes a machine learning-based approach for prioritizing disease-causing variants by integrating genetic data with phenotypic similarity scores. By leveraging deep learning techniques, Xrare enhances the identification of pathogenic variants and adapts to complex genotype-phenotype relationships, making it a powerful tool for clinical diagnostics [7]. AMELIE sets itself apart by leveraging biomedical literature at scale, analyzing millions of PubMed abstracts and full-text articles to support molecular diagnosis. It employs a logistic regression classifier trained on simulated patient data, allowing it to rank causative variants effectively. By dynamically incorporating new scientific findings, AMELIE improves variant interpretation in the evolving landscape of genomic research [8].

InheriNext® is a domain-specific AI-powered tool for the prioritization of genetic variants through two approaches: one utilizes Human Phenotype Ontology (HPO)-based phenotyping [9] and the other focuses on gene panels to identify causative SNPs and INDELs. It employs three scoring systems: phenotype-correlation score, variant pathogenicity score, and disease-similarity score, which aims to offer a more comprehensive framework for analyzing the complexity and diversity of phenotypes.

To evaluate the performance of gene-ranking tools in identifying causative variants, resources from the Global Alliance for Genomics and Health (GA4GH) were adopted as the benchmarking foundation. The GA4GH Phenopacket dataset is particularly well-suited for benchmarking due to its standardized schema, rich phenotypic detail, and expert-curated variant annotations, which together ensure consistency and clinical relevance in evaluation. By leveraging a widely accepted and biologically diverse dataset, this study enables robust and reproducible comparisons across gene-prioritization methods within realistic diagnostic scenarios.

In 2022, GA4GH introduced the Phenopacket Schema, an ISO-approved standard designed to share detailed clinical and genomic information at the individual level. A Phenopacket links phenotypic features with disease diagnoses, patient data, and genetic variants, thereby enabling the construction of accurate disease models [10]. The GA4GH Phenopacket Store (v0.1.21) provides a comprehensive benchmarking dataset comprising of 7,830 Phenopacket samples covering 489 Mendelian and chromosomal diseases linked to 432 genes and 4,263 unique pathogenic alleles. Previous studies have demonstrated the utility of GA4GH Phenopackets as a benchmark for assessing gene-ranking methods [11].

In this study, the ranking performance of InheriNext® is compared against other established diagnostic tools, with the evaluation focusing on the identification of causative gene variants.

## Materials and Methods

To benchmark different variant prioritization approaches, the GA4GH Phenopackets were used to impute a dataset of simulated genetic samples. Genomes from 600 healthy individuals were obtained from the 1,000 Genomes Project, representing genetic diversity across various populations. Considering the population bottlenecks experienced by European, Asian, and American populations—which drastically reduced genetic diversity in these groups [12]—the below method was developed to generate synthetic Whole Exome Sequencing (WES) samples.

Variants from a diverse selection of healthy individuals were pooled to create a rich reservoir from which 50,000 variants were randomly drawn to construct each simulated case of a “healthy” (normal genetic variability) exome. This pooling strategy ensured that the simulated exomes captured a broad spectrum of genetic variability for analysis. Next, data from 7,244 of 7,830 Phenopacket samples that meet the analysis criteria (e.g.: annotated with phenotypic features) were used. Each known disease-causing variant was added into a synthetic healthy exome along with patient’s phenotypic features to simulate a patient with that genetically-driven disease or disorder (Fig1).

**Fig 1.**
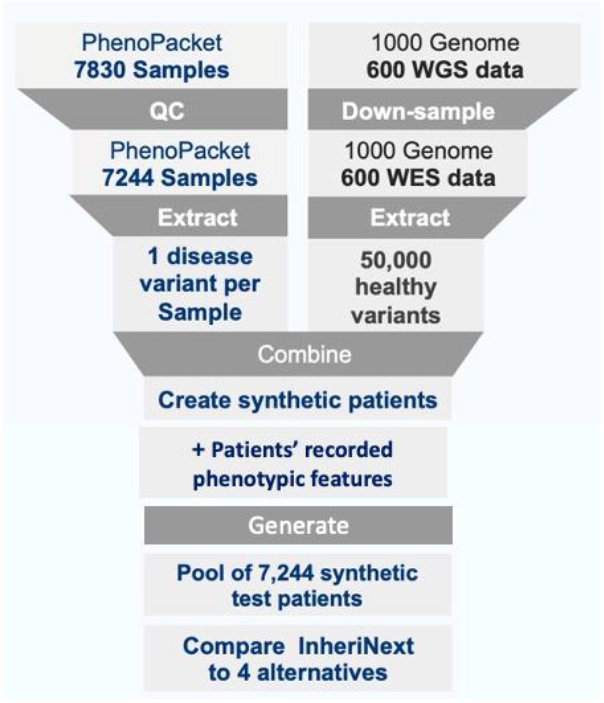
Workflow for generating simulated samples. The diagram outlines the process for generating synthetic patients. Six-hundred (600) healthy individuals’ background genomes were extracted from the 1000 Genome Project, and the causative variants sourced from Phenopacket. After filtering, 7,244 synthetic samples, along with their respective phenotypic features, are created for the following benchmark study.

This process resulted in 7,244 test cases representing disease patients, with each synthesized sample containing annotated phenotypic features, one pathogenic variant from Phenopacket, and alongside 50,000 background variants. These samples were finalized for further analysis and used to evaluate the ranking performance of InheriNext® against four (4) other commonly used variant prioritization tools (A-D).

The benchmark samples encompass nearly five hundred diseases. Grouping similar diseases helps reveal their distribution for the Phenopacket, facilitating a better understanding and allowing for performance analysis across different disease groups. Steps taken to group diseases by K-means clustering using Term Frequency-Inverse Document Frequency (TF-IDF) and Principal Component Analysis (PCA) are described in supplementary data (Fig S1., Fig S2., Fig S3., and Table S1.).

## Results & Discussions

### Ranking Performance in Benchmark Samples

InheriNext® is benchmarked against four (4) different software tools A-D (Exomiser, LIRICAL, Xrare, Amelie) that also utilize phenotype-driven gene prioritization methods to rank candidate pathogenic genes in the 7,244 samples. The “Top-10 Rate” is a practical benchmark for evaluating tool performance, ensuring that the causative gene is captured within the top ranks. It is often visualized using the Cumulative Distribution Function (CDF) plot, which displays the proportion of genes that fall below any given rank. According to previous literature, this method offers an intuitive way to assess and compare the effectiveness of different tools in ranking causative genes [13]. Results show that InheriNext® identified the causative gene within the Top-10 ranks in 98.6% of cases. The corresponding Top-10 rates for tools A, B, C, and D were 95.0%, 86.2%, 87.0%, and 90.9%, respectively (Fig2). The results indicate that InheriNext® achieved the highest Top-10 capture rate in this benchmark comparison.

**Fig 2.**
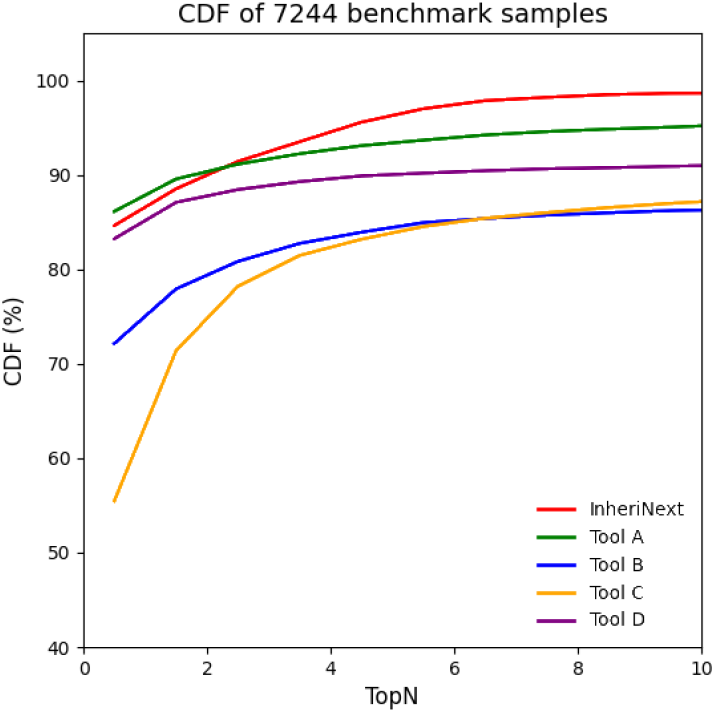
Performance evaluation of InheriNext® with other tools (A-D). The cumulative distribution function (CDF) shows the ranking distribution of causative genes across all five (5) tools. The CDF plots illustrate the percentage of the samples with causative genes ranked within the top*N* by each tool. *N* could be any integer between 1 and 10. Each tool is represented by a different color.

### Ranking Performance in Different Variant Consequences

Variant consequences are pivotal in identifying pathogenic variants, which play a crucial role in diagnosing genetic disorders. The Sequence Ontology (SO) offers standardized annotations to describe the effects of genetic variants on biological sequences, helping categorize their impact on genes, transcripts, and other features to understand their functional implications. (Table S2). This knowledge enables clinicians to pinpoint the genetic basis of a disease accurately, facilitating more precise diagnostics. By assessing the functional impact of these variants, one can evaluate the causative genes more precisely in algorithmic calculations. An analysis of the distribution of variant consequences in the benchmarked samples was conducted, revealing 24 distinct types of variant consequences. In some cases, a variant exhibits multiple types of consequences, resulting in duplicated counts. Table1A indicates that the three (3) most common types of consequences are missense, comprising 49.05% of the total, stop gained at 12.34%, and frameshift truncation accounting for 9.10%. The subsequent evaluation uses the Top-10 rate to assess each tool’s performances in identifying causative genes across different variant consequences (refer to Table1B). InheriNext® achieved the highest Top-10 capture rates for the most frequent, therefore relevant, consequence types (missense variant, stop gained, and frameshift truncation). However, InheriNext® falls short in the synonymous variant category, achieving only a 30.43% in the Top-10 rankings.

### Assessment of Diverse Annotated Phenotype Counts

Phenotypic features, as annotated in the Phenopacket schema, describe clinical symptoms in patients and are used to illustrate the similarity between disease-associated features and those present in a patient. InheriNext® integrates the HPO project [9], which provides an ontology of medically relevant phenotypic features and disease-phenotype annotations, to calculate the likelihood of diseases based on the phenotypic features observed in patients. For example, “arachnodactyly” is a relevant feature for “Marfan syndrome” in HPO database, so if a patient exhibits “arachnodactyly,” they are more likely to have the disease. InheriNext® and other tools use phenotype-driven methods to rank potential pathogenic genes based on phenotypic features in addition to patient genotype data. In the benchmark samples, diverse annotated phenotype counts are inputted for each sample based on phenotypic features in the Phenopacket record used to create that particular simulated case. However, a large count might decrease the performance in ranking causative genes because it may include more unrelated (noise) features, which are not annotated in the disease. For phenotype counts distribution, the results show that: the range of 6-10 phenotypes is most common, seen in 1821 samples, followed closely by the 21-50 range with 1737 samples (Fig3 A). The number of phenotype inputs across each tool’s Top-10s were evaluated to assess performance. The results show that InheriNext® demonstrates consistent performance with percentages ranging from 98% to 100%, indicating strong capability across all phenotype input ranges. Tool A maintains high percentages, reaching 97% in the 6-10 and 11-15 ranges, but drops to 82% in the >50 category. Tool B, Tool C, and Tool D remain steady with over 82% in most ranges; however, they experience a more significant drop in the >50 category (Fig3 B).

**Fig 3.**
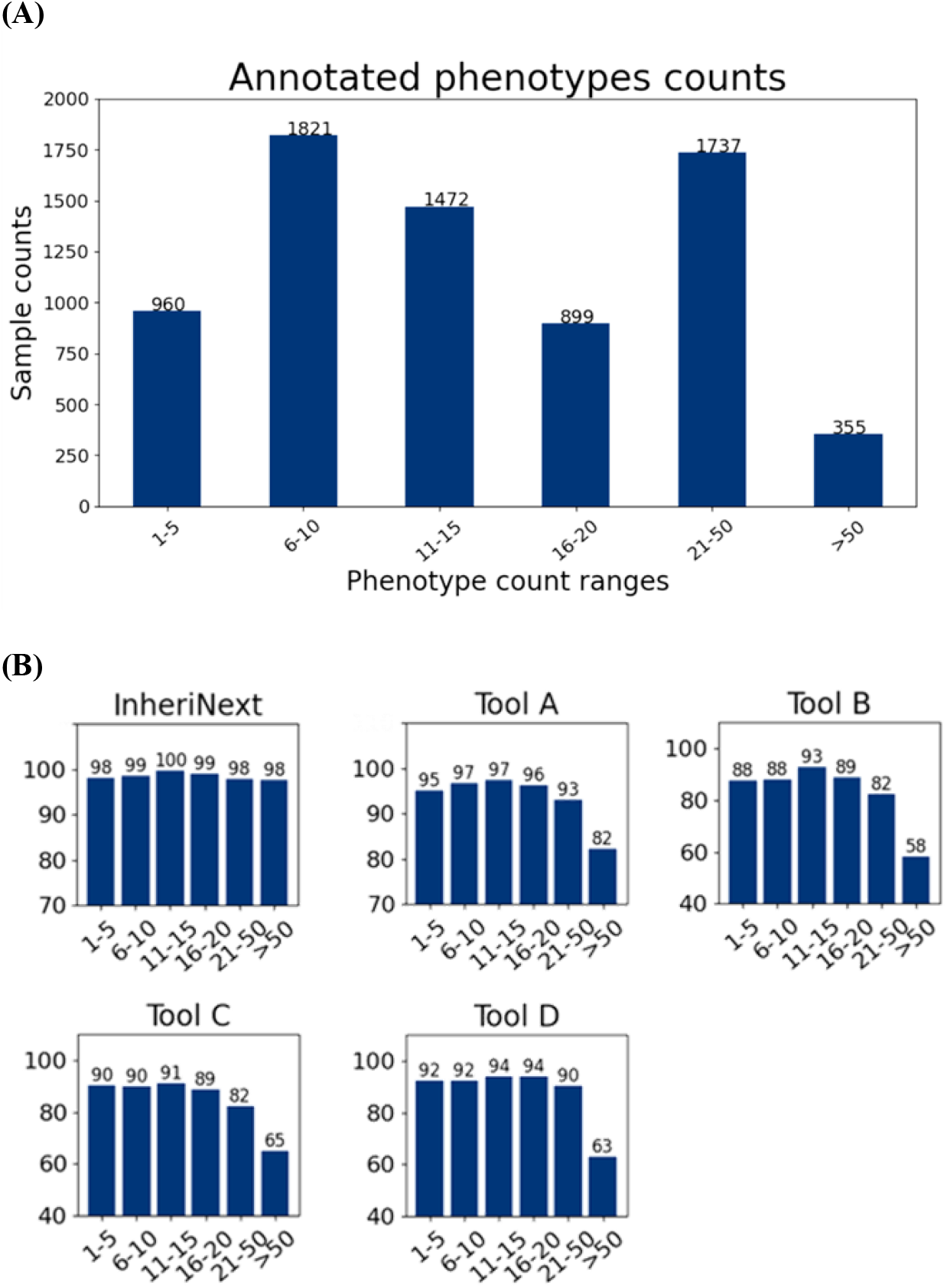
Distribution of Phenotypes Counts in Benchmark Samples. (A) The bar graph illustrates the distribution of samples with different ranges of annotated phenotypic feature counts. Each bar’s height represents the sample count within a specific range. (B) The five bar charts represent the performance of different tools— InheriNext®, Tool A, Tool B, Tool C, and Tool D—across various feature count ranges. Each chart shows the percentage of samples’ causative genes successfully ranked within the Top-10 for each phenotypic feature count range.

### Evaluation of Ranking Performance in Four Disease Groups

Each benchmark sample is annotated with a specific disease. However, some diseases do not match a specific MONDO ID and are therefore removed, leaving 7215 samples for analysis. The Phenopacket-annotated diseases were mapped to second-layer categories from MONDO ontologies (Fig. S1) to generate a word matrix for TF-IDF similarity calculation (Fig. S2 A). The optimal number of clusters for the Phenopacket diseases was determined using k-means clustering with the elbow method applied to the word matrix. This analysis classified the diseases into four major groups, as shown in Fig. S3. To enhance interpretability, these four groups were subsequently renamed with more representative titles, as detailed in Table S1 B.

The scatter plot below displays the distribution of different diseases across four major disease groups, with each data point symbolizing a distinct disease. The size of each point reflects the number of samples for that disease. As reflected by the dense cluster of orange points, this plot clearly shows that the Nervous System and Metabolic System Disorder group has the highest number of samples (Fig4 A). A table is presented that lists examples of actual diseases found in the benchmark samples grouped in four major disease categories (Fig4 B). To evaluate ranking performance, the Top-10 rate was used as an indicator across four disease groups. The results show that InheriNext® consistently ranks over 98% of samples’ causative genes within the Top-10, demonstrating non-discriminatory performance across major disease groups. In comparison, other tools show variability based on the disease type. For example, Tool B performs well in the Cancer or Benign Tumor category but less effectively in the Nervous or Metabolic Disorder group (Fig4 C).

**Fig 4.**
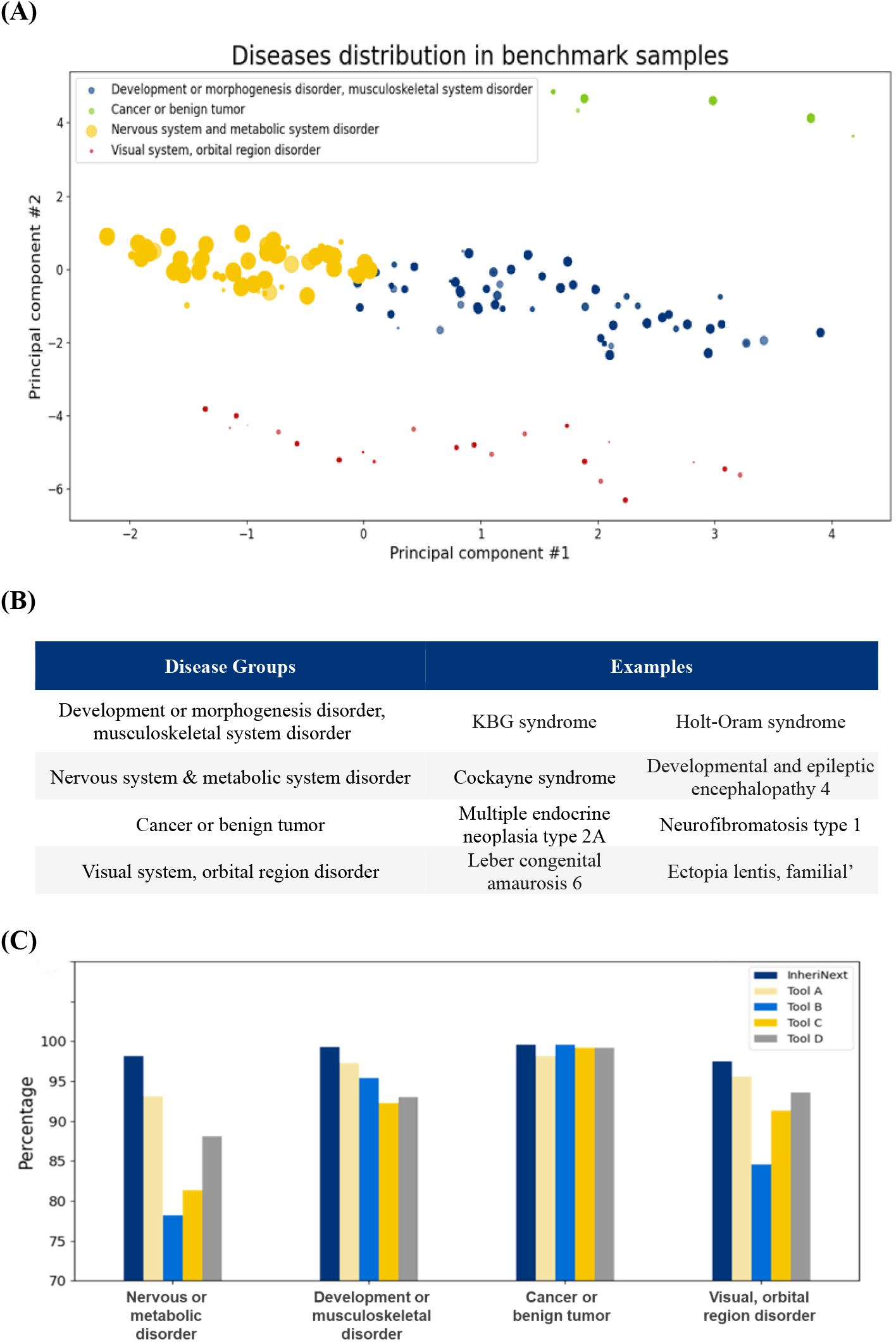
Disease Distribution in Benchmark. (A) The scatter plot illustrates the distribution of diseases by condensing complex data into two principal components (Principal Component #1 and Principal Component #2), highlighting similarities and clustering in four disease groups among the samples. Each point represents a unique disease, and the size indicates the number of samples for that disease; larger points signify a greater number of samples. (B) Example of actual diseases observed in the benchmark samples across four disease groups. (C) The performance comparison across the five tools shows the percentage of samples’ causative genes in Top-10 distributed among the four groups of diseases (abbreviated).

(Note: Disease group names were abbreviated slightly while keeping key terms for clarity and consistency:

1. Development or morphogenesis disorder, musculoskeletal system disorder = Development or musculoskeletal disorder

2. Nervous system and metabolic system disorder = Nervous or metabolic disorder

3. Cancer or benign tumor = Cancer or benign tumor

4. Visual system, orbital region disorder = Visual, orbital region disorder)

Benchmarking tests for InheriNext® showed some key performance strengths, which could be attributed to the following factors:

(1) Integration of Disease Similarity Score Enhances InheriNext® Ranking Performance:

InheriNext® incorporates clinician feedback to refine the prioritization of pathogenic genes. One such enhancement was prompted by a request to improve the assessment of similarity between disease symptoms and patient phenotypes, leading to the integration of a disease similarity score. This feature contributes to stable ranking performance across a variety of disease conditions (Fig4 C). By aligning more closely with clinical realities, this integration supports more accurate and relevant predictions.

(2) Delay Filtering and Removal of Candidates Until After Prioritization: InheriNext® evaluates all potential causative variants and genes, including those with a lower initial likelihood of pathogenicity, to account for their possible clinical relevance. In contrast, other tools may apply early variant consequence filters—for example, excluding 5′ UTR exon variants—which may result in the removal of relevant pathogenic candidates (Table 1B: 5′ UTR exon variant, Tools B–D). InheriNext® retains all candidates through the prioritization stage, enabling more consistent ranking performance across variant types and supporting the identification of atypical or rare variants.

**Table 1.** Distribution of Variant Consequences in Benchmark Samples. (A) The descending proportion and counts of 24 variant consequences identified in benchmark samples. (B) The breakdown of variant consequences by the five (5) tools. The percentages reflect the ability of each tool to rank causative genes within their Top-10s for each consequence.

| Consequences | Proportion % (count) |
| --- | --- |
| Missense variant | 49.05 (4345) |
| Stop gained | 12.34 (1093) |
| Frameshift truncation | 9.10 (806) |
| Coding transcript intron variant | 6.48 (574) |
| Frameshift variant | 4.96 (439) |
| Splice region variant | 4.00 (354) |
| Splice donor variant | 3.04 (269) |
| Frameshift elongation | 2.34 (207) |
| Splice acceptor variant | 1.96 (174) |
| Disruptive inframe deletion | 1.51 (134) |
| Complex substitution | 1.12 (99) |
| Inframe deletion | 0.98 (87) |
| 5 prime UTR exon variant | 0.72 (64) |
| Feature truncation | 0.58 (51) |
| Start lost | 0.52 (46) |
| Multi-nucleotide variant | 0.33 (29) |
| Direct tandem duplication | 0.26 (23) |
| Synonymous variant | 0.26 (23) |
| Disruptive inframe insertion | 0.25 (22) |
| 3 prime UTR intron variant | 0.07 (6) |
| 5 prime UTR intron variant | 0.06 (5) |
| Stop lost | 0.03 (3) |
| Inframe insertion | 0.03 (3) |
| Exon loss variant | 0.03 (3) |

**Table1. Distribution of Variant Consequences in Benchmark Samples.**
| Consequences | InheriNext® | Tool A | Tool B | Tool C | Tool D |
| --- | --- | --- | --- | --- | --- |
| **<u>Missense variant</u>** | **98.64** | 93.9 | 87.59 | 89.07 | 94.38 |
| **<u>Stop gained</u>** | **99.27** | 97.8 | 86.55 | 86.18 | 91.49 |
| **<u>Frameshift truncation</u>** | **99.38** | 98.39 | 91.44 | 89.33 | 93.05 |
| Coding transcript intron variant | **98.08** | 93.55 | 74.22 | 81.88 | 75.44 |
| Frameshift variant | **98.86** | 96.58 | 81.09 | 83.83 | 87.24 |
| Splice region variant | **94.63** | 94.35 | 76.27 | 85.59 | 66.67 |
| Splice donor variant | **99.63** | 98.51 | 86.99 | 86.25 | 95.54 |
| Frameshift elongation | 98.55 | **100** | 92.27 | 92.27 | 96.62 |
| Splice acceptor variant | **99.43** | 94.25 | 86.21 | 83.33 | 91.38 |
| Disruptive inframe deletion | **99.25** | 98.51 | 94.03 | 96.27 | 96.27 |
| Complex substitution | **100** | **100** | 93.94 | 94.95 | 97.98 |
| Inframe deletion | **98.85** | 96.55 | 83.91 | 83.91 | 81.61 |
| 5 prime UTR exon variant | **85.94** | 82.81 | 1.56 | 3.12 | 3.12 |
| Feature truncation | **100** | **100** | **100** | 98.04 | 98.04 |
| Start lost | **100** | 84.78 | 97.83 | 91.3 | **100** |
| Multi-nucleotide variant | **100** | 86.21 | 68.97 | 82.76 | 82.76 |
| Direct tandem duplication | **100** | 86.96 | 73.91 | 95.65 | 86.96 |
| Synonymous variant | 30.43 | 60.87 | 43.48 | **78.26** | 21.74 |
| Disruptive inframe insertion | **100** | 86.36 | 68.18 | 95.45 | 86.36 |
| 3 prime UTR intron variant | **100** | **100** | **100** | **100** | **100** |
| 5 prime UTR intron variant | **100** | 60 | **100** | **100** | **100** |
| Stop lost | 100 | 100 | 33.33 | 100 | 100 |
| Inframe insertion | 100 | 100 | 100 | 100 | 100 |
| Exon loss variant | 100 | 100 | 100 | 100 | 100 |

Despite overall strong performance, several challenges remain. These include limitations in the classification of variants of uncertain significance (VUS) in ClinVar, the underestimation of certain high-allele-frequency variants’ pathogenicity, and ambiguous functional consequences of specific variants. Benchmarking also identified reduced performance in a subset of samples. Further analysis identified two (2) primary contributors, offering clear targets for future refinement:

(1) Variants Annotated With Synonymous Consequence Did Not Rank Well (Table1B):

In its initial implementation, InheriNext® excluded synonymous variants from ranking due to their typically neutral effect relative to non-synonymous variants. This exclusion was intended to streamline prioritization by focusing on more likely pathogenic candidates. However, this approach led to the omission of clinically relevant variants, counter to the broader goal of comprehensive evaluation (see “Delay Filtering and Removal of Candidates Until After Prioritization”). Recent studies have shown that some synonymous variants may have functional consequences, such as impacting splicing motifs or cryptic splice sites, altering microRNA binding [14], and affecting mRNA structure or protein expression via reduced codon optimality [15–17]. In response, InheriNext® has begun to incorporate synonymous variants into the ranking process to improve sensitivity and better reflect their potential clinical relevance.

(2) High Allele Frequency Of The Variants Affect Ranking:

The ClinGen Sequence Variant Interpretation Working Group has refined the ACMG/AMP variant pathogenicity guidelines for rule BA1, which designates variants with a minor allele frequency (MAF) > 0.05 as benign. However, they identified nine (9) variants with MAF > 0.05 that may exhibit pathogenicity. These findings suggest that MAF and pathogenicity do not always correlate. As a result, InheriNext® algorithm may inadvertently penalize genes based on the high allele frequency and affect the ranking accuracy. While high-frequency variants are typically deemed benign, some may still possess pathogenic potential, emphasizing the need to evaluate additional factors when assessing variant pathogenicity. Future optimizations will incorporate additional criteria for adjusting gene scores. The intention is to retain the variant frequency filter as recommended by ACMG/AMP guidelines [18]. However, specific exceptions are systematically reviewed and curated as they appear in the clinical genetics literature, allowing potentially pathogenic variants to be considered despite their relatively high frequency.

## Conclusion

The benchmarking analysis demonstrates that InheriNext® reliably prioritizes candidate pathogenic genes under varied testing scenarios among the tools assessed. In evaluations of phenotype-driven gene prioritization methods, InheriNext® achieved the highest Top-10 capture rate at 98.6% and the lowest missed rate, reflecting high sensitivity. This result is supported by consistent performance across various variant consequences, particularly the most common types—missense variants, stop gains, and frameshift truncations—although some limitations were noted (and addressed) with synonymous variants. Additional tests demonstrate that it maintained ranking accuracy as phenotype complexity increased and across different disease groups, indicating its adaptability across different clinical contexts.

Taken together, these findings suggest that InheriNext® is a reliable tool for gene prioritization, with effective integration of genomic and phenotypic data. Its performance across multiple testing dimensions supports its potential utility to assist in genetic diagnostics and research applications.

## Acknowledgements

The authors acknowledge Melissa Haendel and the Monarch Initiative (https://monarchinitiative.org/) in developing a standardized framework for integrating genomic, environmental, and phenotypic data [10]. Phenopackets provide a critical foundation for ensuring that phenotype profiles are FAIR++ for disease diagnosis and discovery [19]. Their efforts in standardizing data formats, including gene data (GFFs, VCFs, BED files), environmental data, and the novel PXF format, have been instrumental in advancing interoperability and data sharing in rare disease research.

## Availability of Data

The benchmark simulated samples from the GA4GH Phenopacket are available from the corresponding author upon request.

## Notes

### Competing Interest Statement

All authors are either employees or consultants that work or have worked for Compass Bioinformatics, developers of InheriNext.

### Summary of Updates

Updates based on feedback from version 1 has been incorporated.

## References

1. Groft SC, Posada M, Taruscio D: Progress, challenges and global approaches to rare diseases. Acta Paediatr. 2021, 110:2711–2716.

2. Basel-Salmon, L. (2024). Phenotypic compatibility and specificity in genomic variant classification. European Journal of Human Genetics, 32(1), 471–473.

3. Robinson PN, Köhler S, Oellrich A, Sanger Mouse Genetics Project, Wang K, Mungall CJ, Lewis SE, Washington N, Bauer S, Seelow D, Krawitz P, Gilissen C, Haendel M, Smedley D: Improved exome prioritization of disease genes through cross-species phenotype comparison. Genome Res. 2014, 24:340–348.

4. Jacobsen JOB, Kelly C, Cipriani V, Genomics England Research Consortium, Mungall CJ, Reese J, Danis D, Robinson PN, Smedley D: Phenotype-driven approaches to enhance variant prioritization and diagnosis of rare disease. Hum Mutat. 2022 Apr 27;43(8):1071–1081

5. Smedley, D., Jacobsen, J. O. B., Jager, M., Köhler, S., Holtgrewe, M., Schubach, M., Siragusa, E., Zemojtel, T., Buske, O. J., Washington, N. L., Bone, W. P., Haendel, M. A., & Robinson, P. N. (2015). Next-generation diagnostics and disease-gene discovery with the Exomiser. Nature Protocols, 10 (12), 2004–2015. doi:10.1038/nprot.2015.124

6. Robinson, P. N., Ravanmehr, V., Jacobsen, J. O. B., Danis, D., Zhang, X. A., Carmody, L. C., Gargano, M. A., Thaxton, C. L., [UNC Biocuration Core], Karlebach, G., Reese, J., Holtgrewe, M., Köhler, S., McMurry, J. A., Haendel, M. A., & Smedley, D. (2020). Interpretable clinical genomics with a likelihood ratio paradigm. American Journal of Human Genetics, 107 (3), 403–417. 10.1016/j.ajhg.2020.06.021

7. Li, Q., Zhao, K., Bustamante, C. D., Ma, X., & Wong, W. H. (2019). Xrare: A machine learning method jointly modeling phenotypes and genetic evidence for rare disease diagnosis. Genetics in Medicine, 21 (9), 2126–2134. 10.1038/s41436-019-0439-8

8. Birgmeier, J., Haeussler, M., Deisseroth, C. A., Steinberg, E. H., Jagadeesh, K. A., Ratner, A. J., Guturu, H., Wenger, A. M., Diekhans, M. E., Stenson, P. D., Cooper, D. N., Ré, C., Beggs, A. H., Bernstein, J. A., & Bejerano, G. (2020). AMELIE speeds Mendelian diagnosis by matching patient phenotype and genotype to primary literature. Science Translational Medicine, 12 (544). doi:10.1126/scitranslmed.aau9113

9. Robinson P.N., Köhler S., Bauer S., Seelow D., Horn D., Mundlos S.. The Human Phenotype Ontology: a tool for annotating and analyzing human hereditary disease. Am. J. Hum. Genet. 2008; 83:610–615

10. Danis, D., Bamshad, M. J., Bridges, Y., Caballero-Oteyza, A., Cacheiro, P., Carmody, L. C., Chimirri, L., Chong, J. X., Coleman, B., Dalgleish, R., Freeman, P. J., Graefe, A. S. L., Groza, T., Hansen, P., Jacobsen, J. O. B., Klocperk, A., Kusters, M., Ladewig, M. S., Marcello, A. J., Mattina, T., Mungall, C. J., Munoz-Torres, M. C., Reese, J. T., Rehburg, F., Reis, B. C. S., Schuetz, C., Strauss, T., Sundaramurthi, J. C., Thun, S., Wissink, K., Wagstaff, J. F., Zocche, D., Haendel, M. A., & Robinson, P. N. (2025). A corpus of GA4GH Phenopackets: Case-level phenotyping for genomic diagnostics and discovery. Human Genetics and Genomics Advances, 6(1), 100371.

11. Bridges, Y., de Souza, V., Cortes, K. G., Haendel, M., Harris, N. L., Korn, D. R., Marinakis, N. M., Matentzoglu, N., McLaughlin, J. A., Mungall, C. J., Osumi-Sutherland, D., Robinson, P. N., Smedley, D., & Jacobsen, J. O. B. (2024). Towards a standard benchmark for variant and gene prioritisation algorithms: PhEval - Phenotypic inference Evaluation framework. bioRxiv. 2024 Jun 16:2024.06.13.598672.

12. Zheng-Bradley, X., & Flicek, P. (2017). Applications of the 1000 Genomes Project resources. Briefings in Functional Genomics, 16(3), 163–170. 10.1093/bfgp/elw027

13. Masino, A. J., Dechene, E. T., Dulik, M. C., Wilkens, A., Spinner, N. B., Krantz, I. D., Pennington, J. W., Robinson, P. N., & White, P. S. (2014). Clinical phenotype-based gene prioritization: An initial study using semantic similarity and the Human Phenotype Ontology. BMC Bioinformatics, 15, 248. 10.1186/1471-2105-15-248

14. F. Supek, B. Miñana, J. Valcárcel, T. Gabaldón, B. Lehner Synonymous Mutations Frequently Act as Driver Mutations in Human Cancers. Cell, 156 (2014), pp. 1324–1335

15. P. Brest, P. Lapaquette, M. Souidi, K. Lebrigand, A. Cesaro, V. Vouret-Craviari, B. Mari, P. Barbry, J.-F. Mosnier, X. Hébuterne, et al. A synonymous variant in IRGM alters a binding site for miR-196 and causes deregulation of IRGM-dependent xenophagy in Crohn’s diseaseNat. Genet., 43 (2011), pp. 242–245

16. R.A. Bartoszewski, M. Jablonsky, S. Bartoszewska, L. Stevenson, Q. Dai, J. Kap pes, J.F. Collawn, Z. Bebok. A Synonymous Single Nucleotide Polymorphism in ΔF508 CFTR Alters the Secondary Structure of the mRNA and the Expression of the Mutant Protein. J. Biol. Chem., 285 (2010), pp. 28741–28748

17. A. Kim, J. Le Douce, F. Diab, M. Ferovova, C. Dubourg, S. Odent, V. Dupé, V. David, L. Diambra, E. Watrin, M. de Tayrac. Synonymous variants in holoprosencephaly alter codon usage and impact the Sonic Hedgehog protein. Brain, 143 (2020), pp. 2027–2038

18. Kim, S.Y., Kim, B.J., Oh, D.Y. et al. Improving genetic diagnosis by disease-specific, ACMG/AMP variant interpretation guidelines for hearing loss. Sci Rep 12, 12457 (2022). 10.1038/s41598-022-16661-x

19. Haendel, M. (2016). Phenopackets: Making phenotype profiles FAIR++ for disease diagnosis and discovery. figshare. Presentation. 10.6084/m9.figshare.3180898.v1

