## Supplemental Materials for "Evaluating a Standard Benchmark for Gene Prioritization: The InheriNext® Algorithm’s Integration of Genomic and Phenotypic Information"

Supplementary Materials

#### K-means Clustering Diseases Using Term Frequency-inverse Document Frequency (TF-IDF) and Principal Component Analysis (PCA)

##### (1) Building Ontology Structure:

MONDO ontologies are used to standardize disease terminology within a unified framework to establish initial relationships and processes specific MONDO for analysis [1]. The diagram below shows a simplified version of the MONDO ontology (Fig. S1). At the root is "Disease" highlighted in the red box. Following it is "human disease", which branches into several second-layer categories, such as "cardiovascular disorder", "acute disease", "cancer or benign tumor", and "disorder of development or morphogenesis". In MONDO system, these second-layer categories composition terms are broken down into individual words for calculating similarity between diseases.

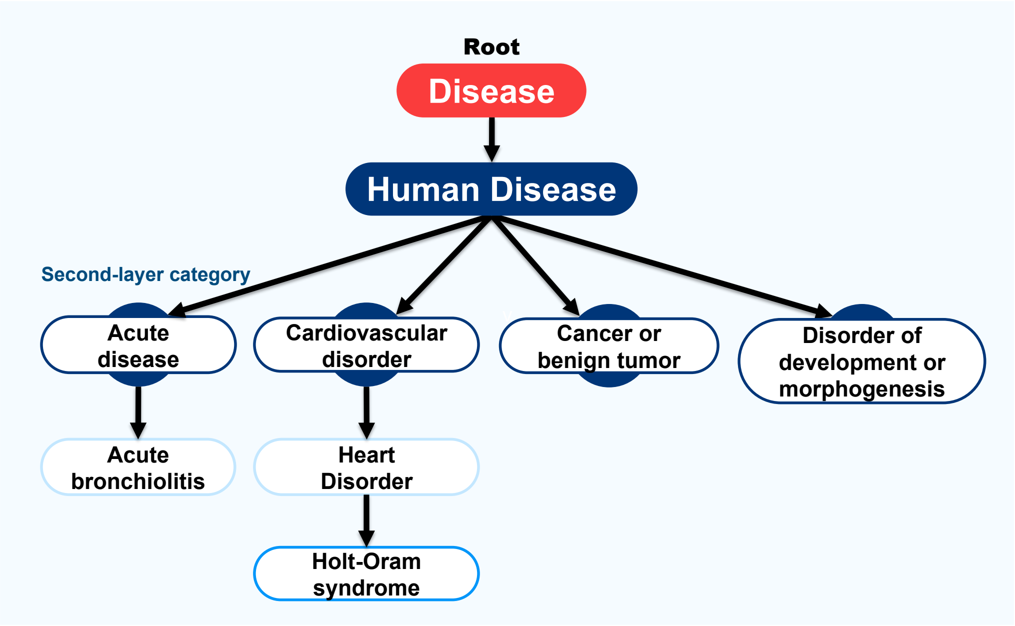

**Fig. S1. The simplified ontology hierarchical structure of diseases in the MONDO System.** This diagram represents a portion of the MONDO disease ontology, illustrating the hierarchical relationships among various diseases. The "Root" node at the center is labeled "Disease," from which several branches extend to different disease categories and sub-categories.

##### (2) Text Processing and Term Frequency-Inverse Document Frequency (TF-IDF) Calculation:

In the Phenopacket-annotated diseases, there are 28 unique second-layer categories from MONDO ontologies. These second-layer categories composition terms are broken down into individual words, with stopwords such as "the", "is", "that" removed, and assembled into a word matrix (Fig. S2 A). Each Phenopacket disease's corresponding second-layer categories are then represented by a set of word vectors, which are converted into TF-IDF vectors for subsequent similarity analysis [2]. The dendrogram was used to interpret how disease terms cluster together based on their TF-IDF similarity scores. In Fig. S2 B, a lower linkage height indicates greater similarity between terms. For example, if two disease terms are joined at a low height, they share a high degree of textual similarity based on their TF-IDF scores. In this analysis, the dendrogram visually represents the hierarchical relationships between disease terms, with the height reflecting their TF-IDF-based distances. It effectively highlights these distances, helping to distinguish closely related disease terms from more distinct ones. Furthermore, dendrograms rely on the ultrametric tree assumption, which rarely holds in real-world text analysis, and may lead to misleading representations of term similarities. To overcome these challenges, Principal component analysis (PCA) and k-means clustering were adopted for disease term classification. PCA reduces the dimensionality of TF-IDF vectors while preserving key features, improving visualization and interpretability. K-means, on the other hand, provides a scalable and flexible clustering approach that determines a specific number of groups, allowing for further evaluation and analysis of these disease clusters.

**(A)**

| | aging | auditory | benign | cancer | cardiovascula | chromosomal | connective | development | digestive | disease | disorder | endocrine | hematologic | immune |
|---|---|---|---|---|---|---|---|---|---|---|---|---|---|---|
| neurodevelopmental disorder with intracranial hemorrhage, seizures, and spasticity | 0 | 0 | 0 | 0 | 0 | 0 | 0 | 0 | 0 | 0 | 1.07494842 | 0 | 0 | 0 |
| hereditary spastic paraplegia 7 | 0 | 0 | 0 | 0 | 0 | 0 | 0 | 1.92668765 | 0 | 3.12934318 | 2.14989684 | 0 | 0 | 0 |
| spermatogenic failure 91 | 0 | 0 | 0 | 0 | 0 | 0 | 0 | 0 | 0 | 0 | 1.07494842 | 0 | 0 | 0 |
| Larsen syndrome | 0 | 0 | 0 | 0 | 0 | 0 | 0 | 1.92668765 | 0 | 0 | 2.14989684 | 0 | 0 | 0 |
| congenital secretory sodium diarrhea 3 | 0 | 0 | 0 | 0 | 0 | 0 | 0 | 0 | 7.1264838 | 0 | 1.07494842 | 0 | 0 | 0 |
| neurodevelopmental disorder with early-onset parkinsonism and behavioral abnormalities | 0 | 0 | 0 | 0 | 0 | 0 | 0 | 0 | 0 | 0 | 1.07494842 | 0 | 0 | 0 |
| muscular dystrophy, limb-girdle, autosomal recessive 28 | 0 | 0 | 0 | 0 | 0 | 0 | 0 | 0 | 0 | 0 | 2.14989684 | 0 | 0 | 0 |
| cleidocranial dysplasia 1 | 0 | 0 | 0 | 0 | 0 | 0 | 0 | 1.92668765 | 0 | 1.56467159 | 2.14989684 | 0 | 0 | 0 |
| platelet-type bleeding disorder 15 | 0 | 0 | 0 | 0 | 0 | 0 | 0 | 0 | 0 | 0 | 1.07494842 | 0 | 4.31779008 | 0 |
| severe combined immunodeficiency, autosomal recessive, T cell-negative, B cell-negative, NK cell-negative, due to adenosine deaminase deficiency | 0 | 0 | 0 | 0 | 0 | 0 | 0 | 0 | 0 | 1.56467159 | 1.07494842 | 0 | 0 | 4.10176399 |
| arthrogryposis, distal, with impaired proprioception and touch | 0 | 0 | 0 | 0 | 0 | 0 | 0 | 1.92668765 | 0 | 0 | 2.14989684 | 0 | 0 | 0 |

**(B)**

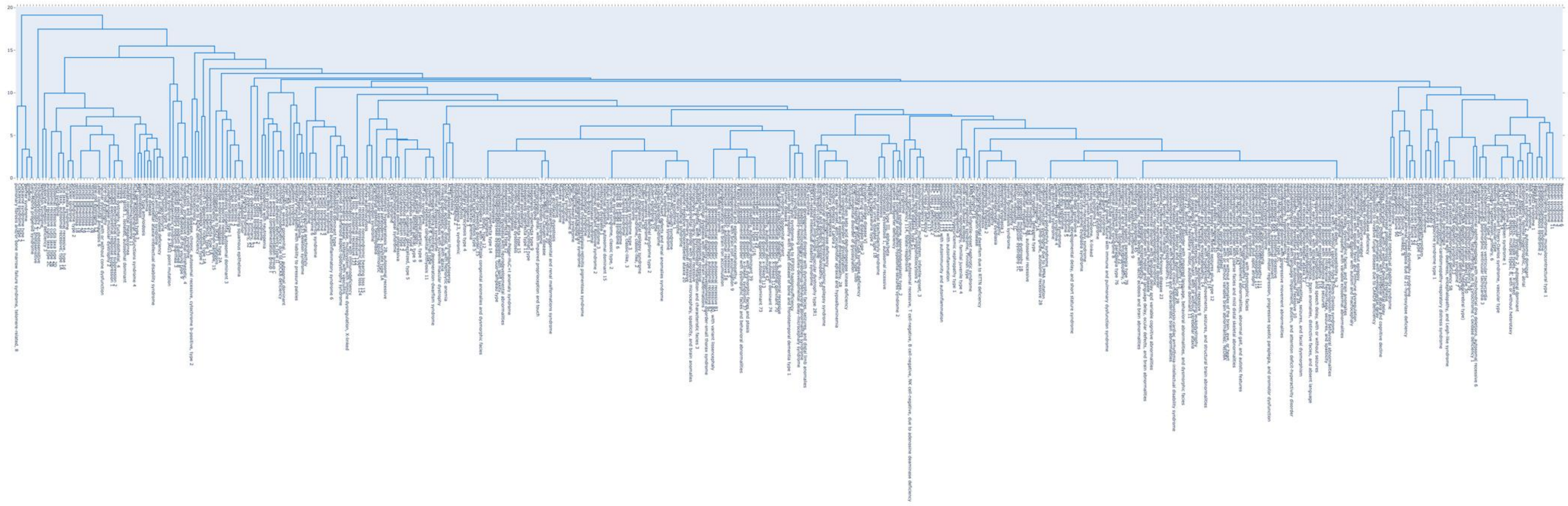

**Fig. S2. The ontology hierarchical relationship among diseases in the dendrogram based on TF-IDF similarity from a word matrix.** (A) A word matrix generated from 28 unique second-layer disease categories, with category names tokenized into individual terms after removing common stop words. (B) The dendrogram illustrates the hierarchical clustering of disease terms based on TF-IDF similarity, where lower linkage heights indicate greater textual similarity. As the hierarchy builds upward, the structure may become less precise in reflecting actual semantic distances.

**(3) Principal Component Analysis (PCA) and K-means Clustering:**
To present the results more intuitively, PCA was applied to reduce dimensionality and capture the most significant differences in the TF-IDF vectors of Phenopacket diseases [3]. This reduction process enabled visualization in a 2D plot. After applying PCA, k-means clustering was used to group the diseases based on these reduced dimensions. K-means clustering is an unsupervised learning algorithm used for grouping unlabeled data points into distinct clusters [4]. To effectively apply k-means, the elbow method [5] was used to determine the optimal number of clusters. The optimal K was identified at the point where Within-Cluster Sum of Squares (WCSS) stops decreasing sharply, indicating a balance between cluster compactness and flexibility. The results showed that when K=4, SSE exhibited a steep decline before leveling off, suggesting that four clusters provided the most effective grouping without overfitting or unnecessary complexity (Fig. S3).

**Fig. S3. The Elbow Method for determining the optimal number of clusters**. K values from one to seven were tested, using WCSS as a performance metric. The plot showed a sharp decline in WCSS when K = 4; beyond this point, adding more clusters did not significantly improve clustering performance, as the rate of WCSS decrease slowed down.

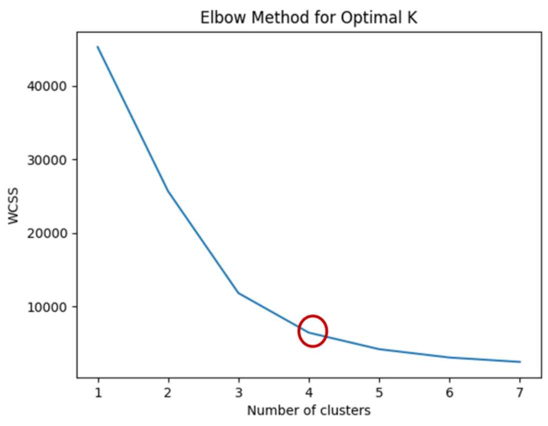

**(4) Defining Representative Disease Terms Across Four Clusters:**
After categorizing the diseases into four groups, the top two highest-frequency terms in each group were selected as representative names by Mondo ontology second-layer disease terms, providing a clearer and more direct overview of diseases in benchmark. When choosing a representative name, the International Classification of Diseases (ICD) was considered, as it offers a comprehensive disease classification system that typically includes standardized disease names [6]. Although *syndromic disease* is a recognized classification in the MONDO ontology system—referring to disorders that

affect multiple organ systems—it cannot be classified under ICD-11 (Table. S1 A). Due to its lack of specificity and limited representative power for precise disease grouping, syndromic disease was excluded to enable a clearer understanding of the disease categories. **Note** that only the terms within each cluster observed in the benchmark are presented in Table S1 A. Finally, representative names were selected from the top two most frequent terms within each cluster to define a more informative and meaningful disease label (Table. S1 B).

**Table S1. Top Two High-Frequency Terms in Each Cluster Defining the Representative Disease Group Name.** (A) ICD-11 is considered for selecting the representative disease term. (B) In the MONDO ontology system, 28 unique second-layer terms are used to classify disease groups, providing a structured hierarchy for disease representation. The table presents the top two highest-frequency terms from each of the four clusters, which will be considered in defining a representative disease term**.**

**(A)**

| ICD-11 for Disease Classifications System | Mondo disease terms (second layer) |
|---|---|
| 01 Certain infectious or parasitic diseases | |
| 02 Neoplasms | cancer or benign tumor |
| 03 Diseases of the blood or blood-forming organs | |
| 04 Diseases of the immune system | |
| 05 Endocrine, nutritional or metabolic diseases | |
| 06 Mental, behavioural or neurodevelopmental disorders | disorder of development or morphogenesis |
| 07 Sleep-wake disorders | |
| 08 Diseases of the nervous system | nervous system disorder |
| 09 Diseases of the visual system | disorder of orbital region<br>disorder of visual system |
| 10 Diseases of the ear or mastoid process | |
| 11 Diseases of the circulatory system | |
| 12 Diseases of the respiratory system | |
| 13 Diseases of the digestive system | |
| 14 Diseases of the skin | |
| 15 Diseases of the musculoskeletal system or connective tissue | musculoskeletal system disorder |

| |
|---|
| 16 Diseases of the genitourinary system |
| 17 Conditions related to sexual health |
| 18 Pregnancy, childbirth or the puerperium |
| 19 Certain conditions originating in the perinatal period |
| 20 Developmental anomalies |
| 21 Symptoms, signs or clinical findings, not elsewhere classified |
| 22 Injury, poisoning or certain other consequences of external causes |
| 23 External causes of morbidity or mortality |
| 24 Factors influencing health status or contact with health services |
| 25 Codes for special purposes |
| 26 Supplementary Chapter Traditional Medicine Conditions |

**(B)**

| Clusters | Mondo disease terms (second layer) | Term frequency | Representative Disease Classification |
|---|---|---|---|
| **Class 0** | disorder of orbital region | 357 | Visual system, orbital region disorder |
| | disorder of visual system | 357 | |
| **Class 1** | cancer or benign tumor | 474 | Cancer or benign tumor |
| | syndromic disease | 474 | |
| **Class 2** | nervous system disorder | 2254 | Nervous system and metabolic system disorder |
| | metabolic disease | 696 | |
| **Class 3** | disorder of development or morphogenesis | 2035 | Development or morphogenesis disorder, musculoskeletal system disorder |
| | syndromic disease | 1925 | |
| | musculoskeletal system disorder | 1185 | |

**Variant Consequence Annotations in the Sequence Ontology (SO)**

The Sequence Ontology (SO) provides a standardized set of annotations for describing the consequences of genetic variants on biological sequences [7]. These annotations help categorize the impact of a variant on genes, transcripts, and other sequence features, facilitating the interpretation of potential functional implications. Each SO term is linked to a unique SO ID and a corresponding description (Table S2). **Note** that only the consequences observed in the benchmark samples are presented.

| SO Term | SO ID | Description |
|---|---|---|
| Missense variant | SO:0001583 | A sequence variant, that changes one or more bases, resulting in a different amino acid sequence but where the length is preserved. |
| Stop gained | SO:0001587 | A sequence variant whereby at least one base of a codon is changed, resulting in a premature stop codon, leading to a shortened polypeptide. |
| Frameshift truncation | SO:0001910 | A frameshift variant that causes the translational reading frame to be shortened relative to the reference feature. |
| Coding transcript intron variant | SO:0001969 | A transcript variant occurring within an intron of a coding transcript. |
| Frameshift variant | SO:0001589 | A sequence variant which causes a disruption of the translational reading frame, because the number of nucleotides inserted or deleted is not a multiple of three. |
| Splice region variant | SO:0001630 | A sequence variant in which a change has occurred within the region of the splice site, either within 1-3 bases of the exon or 3-8 bases of the intron. |
| Splice donor variant | SO:0001575 | A splice variant that changes the 2 base pair region at the 5' end of an intron. |
| Frameshift elongation | SO:0001909 | A frameshift variant that causes the translational reading frame to be extended relative to the reference feature. |
| Splice acceptor variant | SO:0001574 | A splice variant that changes the 2 base region at the 3' end of an intron. |
| Disruptive inframe deletion | SO:0001826 | An inframe decrease in cds length that deletes bases from the coding sequence starting within an existing codon. |

| | | |
|---|---|---|
| Complex substitution | SO:1000005 | When no simple or well defined DNA mutation event describes the observed DNA change, the keyword \"complex\" should be used. Usually there are multiple equally plausible explanations for the change. |
| Inframe deletion | SO:0001822 | An inframe non synonymous variant that deletes bases from the coding sequence. |
| 5 prime UTR exon variant | SO:0002092 | A UTR variant of exonic sequence of the 5' UTR. |
| Feature truncation | SO:0001906 | A sequence variant that causes the reduction of a genomic feature, with regard to the reference sequence. |
| Start lost | SO:0002012 | A codon variant that changes at least one base of the canonical start codon. |
| MNV | SO:0002007 | An MNV is a multiple nucleotide variant (substitution) in which the inserted sequence is the same length as the replaced sequence. |
| Direct tandem duplication | SO:1000039 | A tandem duplication where the individual regions are in the same orientation. |
| Synonymous variant | SO:0001819 | A sequence variant where there is no resulting change to the encoded amino acid. |
| Disruptive inframe insertion | SO:0001824 | An inframe increase in cds length that inserts one or more codons into the coding sequence within an existing codon. |
| 3 prime UTR intron variant | SO:0002090 | A UTR variant of intronic sequence of the 3' UTR. |
| 5 prime UTR intron variant | SO:0002091 | A UTR variant of intronic sequence of the 5' UTR. |
| Stop lost | SO:0001578 | A sequence variant where at least one base of the terminator codon (stop) is changed, resulting in an elongated transcript. |
| Inframe insertion | SO:0001821 | An inframe non synonymous variant that inserts bases into in the coding sequence. |
| Exon loss variant | SO:0001572 | A sequence variant whereby an exon is lost from the transcript. |

**Table S2. Variant Consequence Annotations in the Sequence Ontology.** This table illustrates the standardized annotations provided by the Sequence Ontology to describe the consequences of genetic variants on biological sequences. The columns include the SO ID, the corresponding SO term, and the description of each term. Only the consequences observed in the benchmark samples are presented in this figure.

### References

1.Vasilevsky, N. A., Matentzoglu, N. A., Toro, S., Flack IV, J. E., Hegde, H., Unni, D. R., Haendel, M. A. (2022). Mondo: Unifying diseases for the world, by the world. medRxiv. https://doi.org/10.1101/2022.04.13.22273750

2. Data Mining K-Means Document Clustering using TFIDF and Word Frequency Count, International Journal of Recent Technology and Engineering (IJRTE), vol. 8, no. 2, pp. 2542-2548, July 2019. DOI: 10.35940/ijrte.B1718.078219. License: CC BY-NC-ND 4.0.

3. Ian T. Jolliffe and Jorge Cadima. Principal Component Analysis: A Review and Recent Developments. Philosophical Transactions of the Royal Society A: Mathematical, Physical and Engineering Sciences, vol. 374, no. 2065, April 13, 2016, 20150202. https://doi.org/10.1098/rsta.2015.0202.

4. Chris Ding and Xiaofeng He. 2004. K-means clustering via principal component analysis. In Proceedings of the twenty-first international conference on Machine learning (ICML '04). Association for Computing Machinery, New York, NY, USA, 29. https://doi.org/10.1145/1015330.1015408

5. M. A. Syakur et al. Integration K-Means Clustering Method and Elbow Method for Identification of the Best Customer Profile Cluster. IOP Conference Series: Materials Science and Engineering, vol. 336, The 2nd International Conference on Vocational Education and Electrical Engineering (ICVEE), November 9, 2017, Surabaya, Indonesia. 2018. IOP Conf. Ser.: Mater. Sci. Eng. 336 012017. DOI: 10.1088/1757-899X/336/1/012017.

6. World Health Organization. (2019). QE84 Acute stress reaction. In *International statistical classification of diseases and related health problems* (11th ed.). https://icd.who.int/browse11/l-m/en#http%3a%2f%2fid.who.int%2ficd%2fentity%2f505909942

7. Eilbeck, K., Lewis, S. E., Mungall, C. J., Yandell, M., Stein, L., Durbin, R., & Ashburner, M. (2005). The Sequence Ontology: a tool for the unification of genome annotations. *Genome Biology, 6*(8), R44. https://doi.org/10.1186/gb-2005-6-8-r44
